# Horizontal transfer of a 180-kbp genomic fraction among the largest viral genomes

**DOI:** 10.1101/2025.04.28.650931

**Authors:** Hiroyuki Hikida, Ruixuan Zhang, Jingjie Chen, Yusuke Okazaki, Hiroyuki Ogata

## Abstract

Viruses are generally considered tiny biological entities with small genomes; however, some dsDNA viruses, known as giant viruses, have large genomes that are comparable to those of small bacteria. Previous studies indicated that the genomes of giant viruses expanded from a small ancestor by a combination of gene duplication, *de novo* gene creation, and horizontal gene transfer (HGT). Among them, virus-to-virus HGTs are recently recognized as an important mechanism for disseminating functional genes between giant viruses. In this study, we isolated a giant virus of a member of pandoraviruses, which have the largest genome sizes reaching 2.5 Mbp. Based on the average nucleotide identity of known pandoraviruses, this new pandoravirus belongs to an existing viral species. However, its genome was approximately 200 kbp larger than those of the other strains from the same species. A genome-wide comparison identified a 180-kbp region with 168 genes missing from the other strains but found in phylogenetically distant pandoraviruses. The GC ratio and homology of the deduced amino acid sequences in this 180-kbp region suggest that the new virus horizontally acquired this region from a distantly related pandoravirus. The gene composition in the 180-kbp region further indicates that this region was already large at the time of the horizontal transfer. Our findings suggest that pandoraviruses can horizontally exchange a large portion of their genomes. This event presumably represents one mechanism for accelerating genomic evolution and gigantism in giant viruses.

## Introduction

Viruses were considered to be tiny biological agents until the discovery of giant viruses that have large particles and genome sizes comparable to those of small bacteria (La Scola et al. 2003; Raoult et al. 2004; Lwoff 1957). These giant viruses primarily infect unicellular eukaryotes and are currently classified into the viral phylum *Nucleocytoviricota* along with some animal viruses (e.g., poxviruses) (Aylward et al. 2021). Recent studies discovered giant viruses in large metagenomic data and revealed their vast diversity and broad distribution in various environments (Endo et al. 2020; Schulz et al. 2020). In these environments, giant viruses modulate nutrient cycles not only by killing the hosts but also by reprogramming host metabolism using a large repertoire of genes related to cellular machinery (Moniruzzaman et al. 2020a; Rosenwasser et al. 2016; Suttle 2007).

Previous studies indicated that giant viruses have evolved from a small ancestor (Iyer et al. 2006; Yutin et al. 2014), acquiring genes by a combination of gene duplication (Suhre 2005; Machado et al. 2023), *de novo* gene creation (Legendre et al. 2018; Forterre and Gaïa 2016), and horizontal gene transfer (HGT) (Irwin et al. 2022; Wu et al. 2024). HGT has a prominent role in expanding the viral gene repertoire and increasing viral fitness, as known for cellular organisms (Keeling 2009; Soucy et al. 2015). Giant viruses have experienced a massive number of HGTs from host organisms and acquired genes related to cellular functions (Kijima et al. 2024; Irwin et al. 2022). At the same time, giant viruses contribute to the evolution of host eukaryotes by providing functional genes that evolved in viral genomes (Da Cunha et al. 2022; Guglielmini et al. 2019). In addition to these host-virus interactions, HGTs among viruses have played a pivotal role in giant virus evolution to disseminate genes acquired from hosts within the viruses (Wu et al. 2024; Kijima et al. 2024).

Pandoraviruses are one of the largest members of the phylum *Nucleocytoviricota* with 1-µm amphora-shaped virions and 1.5–2.5 Mbp genomes (Legendre et al. 2018; Philippe et al. 2013; Aherfi et al. 2018). Their genomes encode 1,000–2,000 genes, most of which are functionally uncharacterized proteins (Abergel et al. 2015). Similar to other nucleocytoviruses, pandoraviruses encode genes involved in cellular functions and viral replication; however, they lack some hallmark genes (e.g., major capsid proteins) (Aherfi et al. 2022; Philippe et al. 2013). Phylogenetically, pandoraviruses are related to phycodnaviruses that infect eukaryotic algae but are highly divergent in terms of genome sizes and morphology (Yutin and Koonin 2013; Aylward et al. 2021). A previous study suggested that another virion protein was substituted for the major capsid proteins, allowing their distinct morphology and genome sizes (Krupovic et al. 2020).

In this study, we isolated a new pandoravirus strain from freshwater lake sediment. This virus was classified into an existing species; however, its genome was 200 kbp larger than other strains of the species. Genome-wide comparisons identified a 180-kbp region with 168 genes, which is missing from the closely related viruses but found in distantly related viruses. Further analysis suggested that this region was horizontally transferred from another virus. Our results suggest that the pandoravirus genomes are highly flexible and can horizontally transfer large genomic factions, which accelerate genomic evolution in giant viruses.

## Results

### Isolation and phylogenetic analysis of a pandoravirus

We isolated a virus from a sediment sample of a Japanese freshwater lake (Lake Biwa) by a co-culture method using a free-living amoeba, *Acanthamoeba castellanii*. The isolated virus had amphora-shaped virions (∼1 µm), which were morphologically similar to that of pandoraviruses (Fig. 1A). The whole genome sequences were determined by a hybrid assembly of short- and long-read sequences, which yielded a 1,970,084-bp contiguous genomic sequence of the virus (Fig. 1B). The newly isolated viruses exhibited 97.2% and 98.3% average nucleotide identity (ANI) to pandoravirus japonicus (PanV-jap) (Hosokawa et al. 2021) and pandoravirus pampulha strain 8.5 (PanV-pam) (Pereira Andrade et al. 2019), respectively. Two tRNA and 1,777 protein-coding genes were predicted. Similar to other pandoaviruses, most of these protein-coding genes were homologous to pandoravirus proteins that are unique to these viruses or ORFans (Fig. 1B and Table S1). Phylogenetically, pandoraviruses are divided into two clades (i.e., A and B) (Legendre et al. 2018). We found that Clade A can be further divided into two subclades (A-I and A-II). The nucleotide identity indicated that the newly isolated virus belongs to Clade A-I (Fig. 1C). A phylogenetic tree based on the concatenated genes conserved in the phylum *Nucleocytovirivota* (i.e., AA18 helicase and RNA polymerase beta subunit 1) confirmed that the newly isolated virus is closely related to PanV-jap and PanV-pam in Clade A-I (Fig. 1D). Current frameworks for giant virus taxonomy often use 95% ANI as a species boundary (Aylward et al. 2021). On the basis of ANI-based clustering and gene trees, PanV-jap, PanV-pam, and the newly isolated virus appeared to form a species. Thus, we considered the newly isolated virus a new strain of *Pandoravirus pampulha* (the name of the first isolate among these three viruses) and designated it pandoravirus pampulha Biwa (PanV-biw).

**Figure 1.**
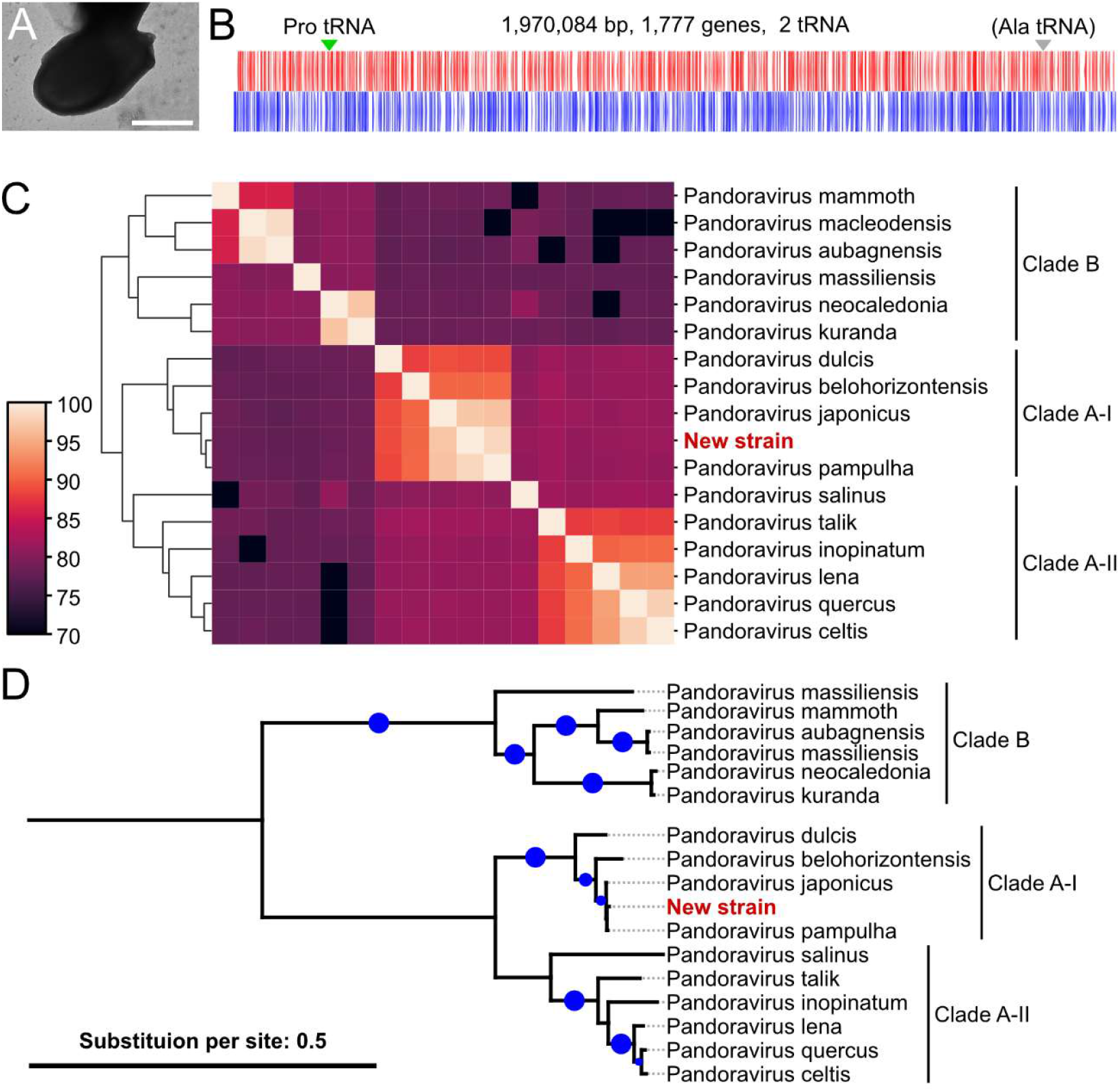
Isolation and phylogenetic analysis of a pandoravirus. (A) Negative stain image of a pandoravirus newly isolated from a Japanese freshwater lake, Lake Biwa (PanV-biw). Bar = 500 nm. (B) Genomic structure of PanV-biw. Red and blue indicate genes located on the leading or complementary strands, respectively. The locations of tRNA genes are indicated by arrowheads. The tRNA^Ala^ gene was predicted as a pseudogene. (C) Comparison of the average nucleotide identity (ANI) between pandoravirus genomes. Viruses were clustered based on pairwise ANI. The order in the columns is the same as that in the rows. (D) A concatenated tree of AA18 helicase and RNA polymerase B subunit 1. The substitution model was Q.plant+F+I+G4. Blue dots indicate the branch that was statistically supported (SH-aLRT >80 and UFB >95).

### PanV-biw contains a 180-kbp region missing from its close relatives but found in its distant relatives

Despite the close phylogenetic relationship, PanV-biw has a genome approximately 200 kbp larger than that of PanV-jap (1,798,487 bp) and PanV-pam (1,676,110 bp). This suggests rapid genome expansion or a reduction among these three viruses (Fig. S1). Genome-wide comparisons between pairs of the pandoraviruses identified a 180-kbp region encoding 168 genes in the PanV-biw genome, which is missing from PanV-pam and PanV-jap (Figs. 2A and S2A). This region was located from 1,107,212 to 1,286,311 nt of PanV-biw and was missing in genomes of other Clade A-I viruses (Fig. S2A).

**Figure 2.**
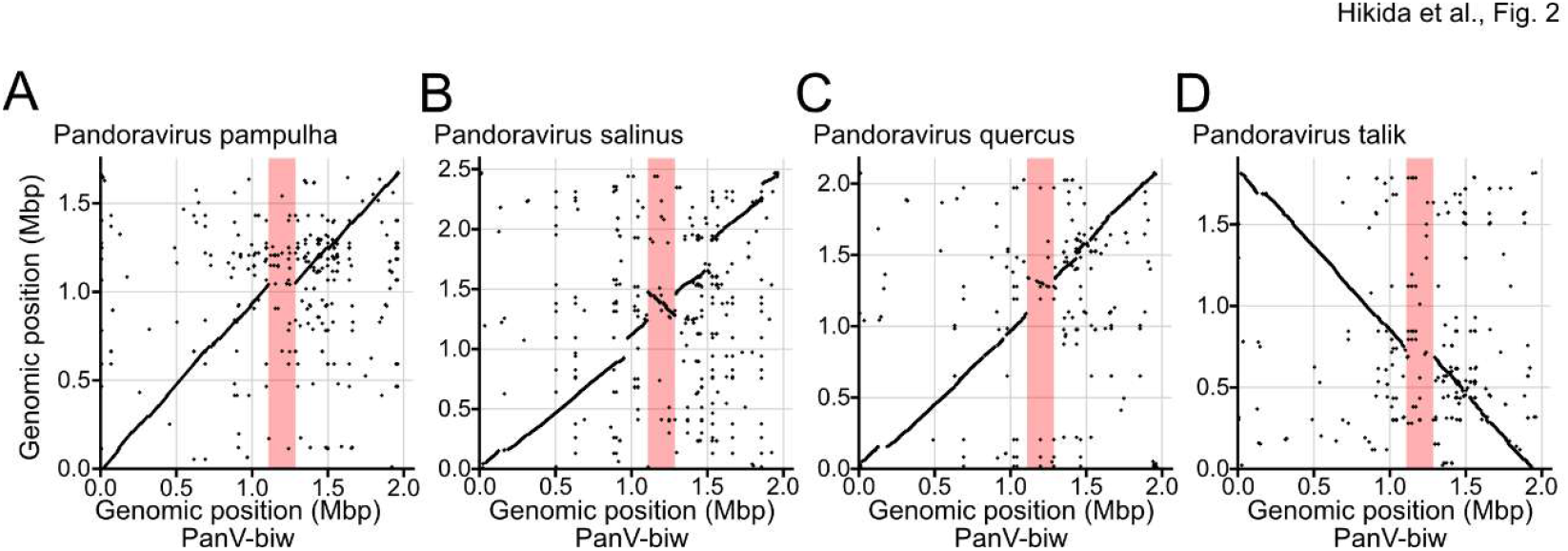
The 180-kbp region missing from close relatives of PanV-biw. Genomic comparison between PanV-biw and (A) pandoraviruses belonging to Clade A-I and (B– D) those belonging to Clade A-II based on BLASTN. X- and Y-axes represent the genomic position in PanV-biw and other pandoraviruses, respectively. The viruses are designated at the top left. (A–D) Red boxes indicate the 180-kbp region missing from other Clade A-I viruses.

To examine the possibility that this 180-kbp region resulted from any artifact in the assembly process, we mapped the raw sequence reads that were used to assemble PanV-jap to the PanV-biw genome. Because these two viruses exhibit high nucleotide identity, most reads were mapped to the PanV-biw genome. However, the 180-kbp region was not covered by the reads from PanV-jap (Fig. S3A). Some PanV-jap reads bridged the 5′ and 3′ flanking regions of the 180-kbp region, confirming the absence of the 180-kbp region in PanV-jap (Fig. S3B). We also mapped the raw long reads from PanV-biw to the PanV-biw genome and identified reads covering the junctions between the 180-kbp and flanking regions (Fig. S3C, D). These results exclude the possibility of artifacts in the assembly process of the genomes.

Further genome-wide comparisons between PanV-biw and other pandoraviruses identified regions homologous to this 180-kbp region (hereafter referred to as “the homologous region”) in phylogenetically distant pandoraviruses (Figs. 2B and S2B). Two viruses from Clade A-II, pandoravirus salinus (PanV-sal) and pandoravirus inopinatum, conserved almost the entire region, although their homologous regions were inverted. Other Clade A-II viruses (pandoravirus quercus and pandoravirus celtis) conserved a part of the region (Figs. 2C and S2C). In the genome of another Clade A-II virus, pandoravirus talik, the region was missing as in the Clade A-I viruses (Fig. 2D). A comparison with the Clade B viruses revealed that the 5′ half flanking region of the 180-kbp region was relatively well conserved, whereas the 3′ half flanking region was not conserved in the Clade B viruses (Fig. S2D), suggesting that this 180-kbp region was located at the boundary of the bipartite structure of the pandoravirus genome described in a previous study (Legendre et al. 2018). These results propose two major possibilities: this region was present in the common ancestor of the Clade A viruses but was lost in most Clade A-I viruses and pandoravirus talik, or the region was horizontally transferred from a Clade A-II virus to PanV-biw.

### The protein and nucleotide similarity suggests a horizontal transfer of the 180-kbp region

The proteins encoded in the 180-kbp region of PanV-biw showed higher identity to those in PanV-sal than the proteins encoded in the remainder of the PanV-biw genome did (Fig. 3A, C). This result suggests that the 180-kb region of PanV-biw has different evolutionary origins from the remainder of the genome. We also examined the genomic GC ratio, which is associated with each lineage of pandoraviruses (Fig. S1). The GC ratio of the 180-kb region was 59.8%, which was lower than 63.5% for the remainder of the PanV-biw genome (Fig. 3B, D). The ratio was also lower than 61.7% of the entire genome of PanV-sal (Fig. 3D). However, the homologous region in PanV-sal showed 59.6% GC, which was close to the ratio of the 180-kbp region of PanV-biw (Fig. 3D). This suggests a shared evolutionary origin of the 180-kbp region of PanV-biw and its homologous region of PanV-sal. These results support the horizontal transfer of the 180-kbp region to PanV-biw rather than the vertical inheritance in Clade A-I. Of note, the deviation of the GC ratio in the homologous region in PanV-sal implies that the region in PanV-sal was also horizontally acquired. Thus, an alternative scenario would be that the 180-kbp region and its homologous region both originated from an unknown entity.

**Figure 3.**
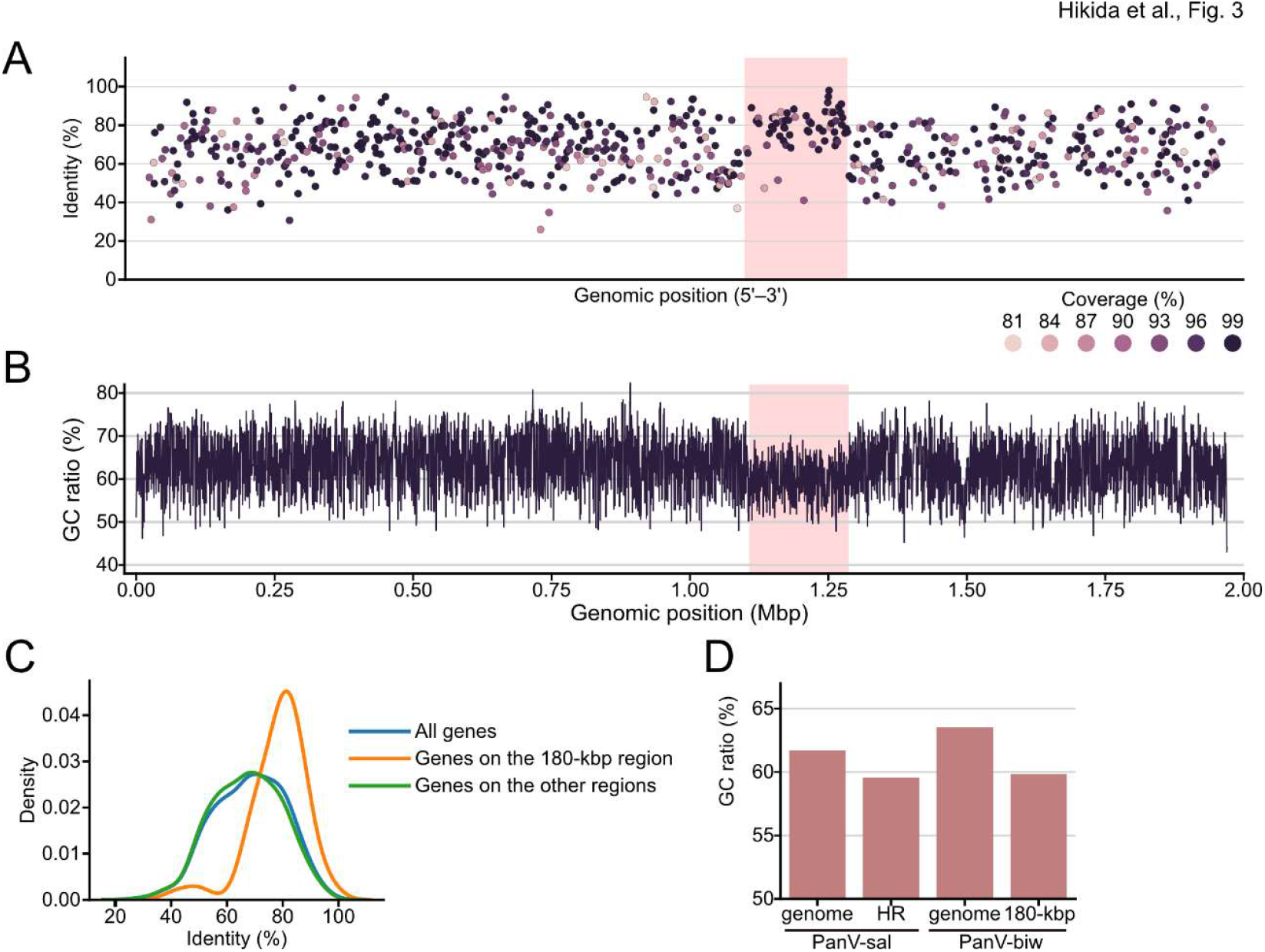
The protein similarity and GC ratio suggest the horizontal transfer of the 180-kbp region to PanV-biw. (A) Identity of proteins encoded in the PanV-biw genome compared with those encoded in the pandoravirus salinus (PanV-sal) genome. The color indicates the coverage of the BLAST alignment. The position of the genes in PanV-sal is shown in Figure S4. (B) The GC ratio within a 500-bp window in the genome of PanV-biw. (A, B) Red boxes indicate the 180-kbp region of PanV-biw. (C) Kernel-density estimate for the distribution of the identity of proteins encoded in the PanV-biw genome to those encoded in the PanV-sal genome. Blue, orange, and green indicate the proteins encoded in the entire genome, the 180-kbp region, and the remainder of the genome, respectively. (D) The GC ratio of PanV-biw and PanV-sal. GC ratio in the entire genome and that within the 180-kbp region (180-kbp) of the PanV-biw genome or the homologous region of the PanV-sal genome (HR) are shown.

### The gene composition of the 180-kbp region suggests genome expansion before the horizontal transfer

Although our results support that the 180-kbp region of PanV-biw was horizontally acquired, it is unclear whether this region was already large at the time of the horizontal transfer or expanded from a small fragment after the transfer. To infer whether the 180-kbp region of PanV-biw was expanded before or after the horizontal transfer to PanV-biw, we examined the gene composition of the 180-kbp region based on the orthologous groups (OGs) shared by the pandoraviruses. Of the 168 genes encoded in the 180-kbp region, 119 genes were assigned to 83 OGs (Fig. 4A). Twenty OGs have more than one copy of the genes in the 180-kbp region of the PanV-biw genome, suggesting that the gene duplication contributed to the expansion of this region. In the phylogenetic tree of a highly duplicated OG (i.e., one of the proteins with the morn-repeat domain), the genes encoded in the 180-kbp region exhibited a phylogenetically distant relationship, which suggests ancient duplication of the OG before the horizontal transfer (Fig. 4B). The genes encoded in the 180-kbp region were grouped with the PanV-sal or PanV-ino genes. These groupings were statistically supported, indicating that they were duplicated in the common ancestral region of these pandoraviruses (Fig. 4B). Of the other 63 OGs encoded in the 180-kbp region of PanV-biw, 56 have homologs in other homologous regions of the Clade A-II pandoraviruses (Fig. 4A). Most of the genes are conserved in PanV-biw, PanV-sal, and PanV-ino, suggesting that their ancestral region already contained a substantial gene repertoire. Collectively, our results suggest that the 180-kbp region was already expanded before the horizontal transfer to PanV-biw with a combination of gene duplication and other mechanisms.

**Figure 4.**
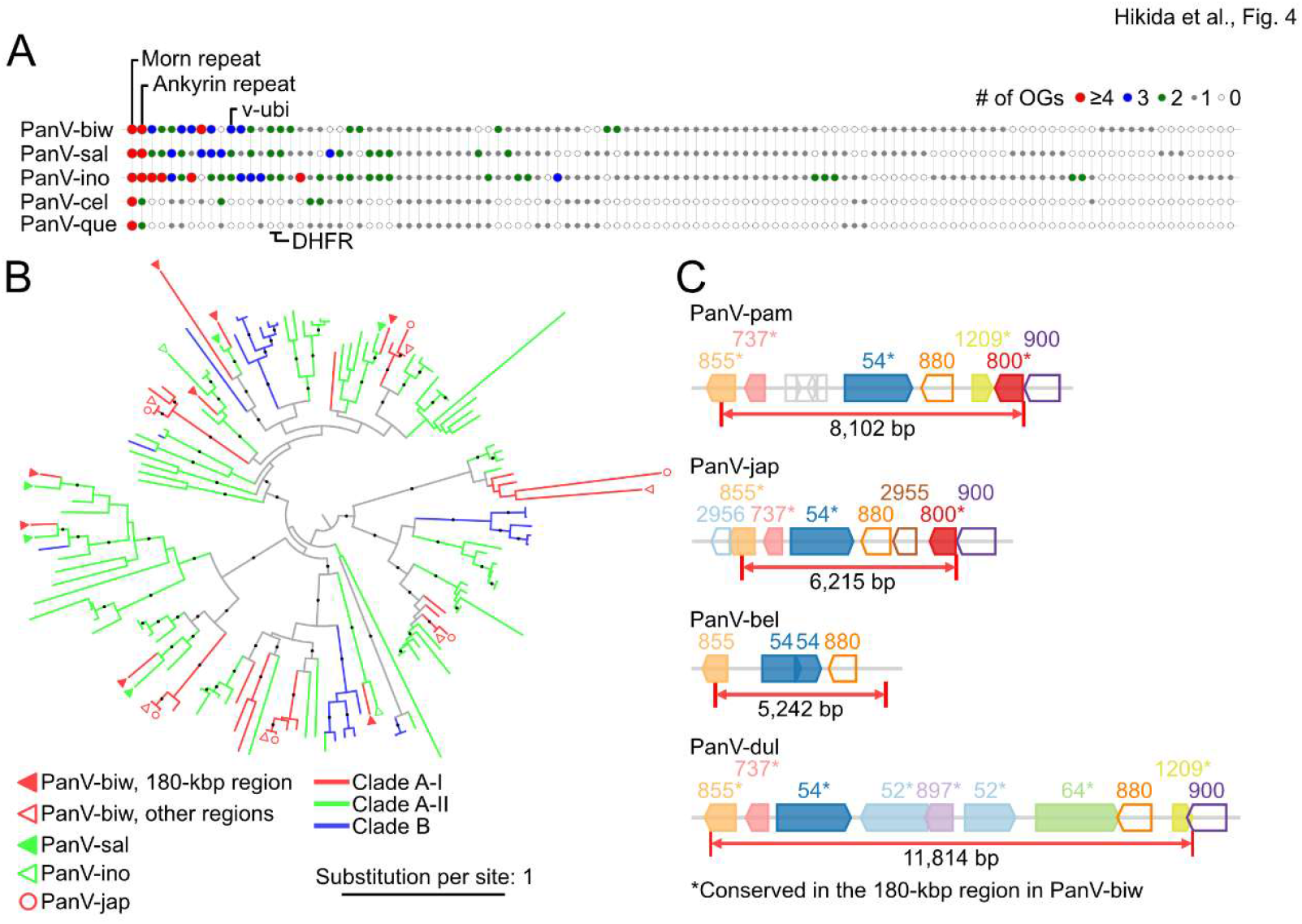
Genome expansions before horizontal transfer. (A) Orthologous groups (OGs) of genes encoded in the 180-kbp region of PanV-biw and its homologous regions in the Clade A-II pandoraviruses. Rows and columns represent genomes and OGs, respectively. Colors indicate the number of genes belonging to each OG, and empty circles indicate that no OGs are encoded in the genome. OGs are ordered by the total number of genes assigned to each OG. An annotation was provided for some OGs. DHFR and v-ubi indicate the dihydrofolate reductase incomplete domain-containing protein and viral ubiquitin, respectively. (B) A phylogenetic tree of the OG of morn-repeat domain proteins. Branch colors indicate the viral clades. The shape and color of the outer nodes represent the origins of the sequences. (C) Genome structure of the corresponding regions in the Clade A-I pandoraviruses. Each box represents genes with direction in the genome. The same numbers and colors indicate genes belonging to the same OGs. Genes homologous to those encoded in the 180-kbp region are marked by an asterisk and shown as filled boxes.

### Clade A-I pandoraviruses have a region missing in PanV-biw

To infer possible mechanisms underlying the horizontal transfer, we investigated the flanking ends of the 180-kbp region of PanV-biw and their corresponding regions in the other Clade A-I pandoraviruses. Although the other Clade A-I pandoraviruses appeared to lack the corresponding region to the 180-kbp region in the dot plots (Figs. 2 and S2), enlarged dot plots identified several kilobase regions in the genomes of Clade A-I viruses that were missing from PanV-biw (hereafter referred to as “the corresponding region”) (Fig. S5). This result suggests that the acquisition of the 180-kbp region was not an insertion but a replacement of the corresponding region by the 180-kbp region. Because a previous study showed that pandoraviruses are capable of recombining their genomic DNA with exogenous elements (Bisio et al. 2023), the putative genomic replacement likely occurred by homologous recombination.

The corresponding regions encoded genes homologous to those encoded in the 180-kbp region of PanV-biw (Fig. 4C). This result suggests that the 180-kbp region of PanV-biw, its homologous region in the Clade A-II pandoraviruses, and its corresponding regions in the Clade A-I pandoraviruses are evolutionarily related. The length of the corresponding regions varied from ∼5 kb in PanV-bel to ∼11 kb in PanV-dul. PanV-dul encoded duplicated genes and more homologous genes to those in the 180-kbp region of PanV-biw than the other Clade A-I viruses. These results indicate that the Clade A-I viruses also experienced expansion and reduction in this region.

### The function of the 180-kbp region is elusive

In the 180-kbp region, most genes encode hypothetical proteins, repeat-containing proteins (i.e., ankyrin-repeat proteins), and genes annotated by their domain structures (i.e., F-box, morn-repeat, and ring domain). Only a few genes had functional annotations. One of these genes contains a dihydrofolate reductase (DHFR) incomplete domain-containing protein (Fig. 4A). DHFR is involved in the synthesis of thymidine (Blaney et al. 1984), which might be beneficial for maintaining the slightly AT-rich sequences of the 180-kbp region. Another example is viral ubiquitin (Fig. 4A), but its function is currently unclear in giant viruses.

## Discussion

In the present study, we isolated a pandoravirus from freshwater lake sediment. This virus represented a new strain of an existing viral species but harbored a 180-kbp region missing from other closely related viruses. The genomic comparison suggests a scenario in which a large genomic fraction was horizontally transferred to the new pandoravirus. Our results suggested that giant viruses are capable of transferring large genomic fractions to other viruses horizontally. Recent studies identified large integrations of giant virus genomes into host chromosomes, which facilitate the transfer of a large proportion of genes from viruses to their hosts (Moniruzzaman et al. 2020b; Zhao et al. 2023). Our results indicated that such large-scale HGTs were also possible among viruses, which may have contributed to their genomic evolution.

This study revealed that closely related pandoravirus strains within the same species exhibited a 200-kbp difference in their genome size, accounting for approximately 10% of their genome size. This finding highlights the high flexibility of the pandoravirus genomes. Pandoraviruses have lost the double-jerry role-type major capsid proteins, which are typical of nucleocytoviruses. Instead, they use other proteins to form virions. This morphological innovation is a potential driver for the development of large virions and genomes (Philippe et al. 2013; Krupovic et al. 2020; Bisio et al. 2023). However, the resulting large genomes apparently contain a lot of unnecessary components and are invaded by mobile genetic elements, such as introns, inteins, and transposable elements (Philippe et al. 2013; Akashi and Takemura 2019; Sun et al. 2015). An experimental study showed that a 600-kbp region is dispensable for viral replication, supporting the large capacity of pandoraviruses to encode nonessential genes (Bisio et al. 2023). Consistent with these previous findings, our discovery of a 180-kbp horizontally transferred region supports that the pandoravirus genomes have high flexibility, which may increase their coding capacity.

Currently, the precise evolutionary trajectory of this region is unknown. Its related regions were found in a wide range of the Clade A pandoraviruses, but their size and gene content are highly diverse. These results suggest frequent genomic expansions and reductions in these regions. Moreover, the GC ratio of the homologous region in PanV-sal implies that these regions have mobility. Pandoraviruses have bipartite genomic structures, which separate the region encoding the core genes and that encoding accessory ones (Legendre et al. 2018). The 180-kbp region of PanV-biw and its homologous or corresponding regions are closely located at the junction of the bipartite structures. Therefore, this region might have distinct properties from the remainder of viral genomes and particularly high flexibility. Recently, the diversity of giant viruses was largely expanded by metagenomic studies. However, metagenomic approaches often miss this type of region because of technical challenges associated with intra-species diversity. Our study indicates that isolation studies are still important to provide a broader overview of the genomic evolution in giant viruses.

The ecological or biological benefits of these regions remain elusive because of limited functional information. One small clue is the DHFR homologs found in the 180-kbp region, which appear beneficial for this region rather than the entire genome. These genes indicate that this region might have selfish properties, although additional functional analyses are required. Recently, a reverse-genetics system for pandoravirus was developed (Bisio et al. 2023; Philippe et al. 2024). Further studies with such experimental systems may reveal the functional importance of these regions and provide novel insights into the role of virus-to-virus HGT in giant virus evolution.

## Methods Virus isolation

Virus isolation and purification were performed as described previously (Hikida et al. 2023). Briefly, a top layer of sediment was previously collected from the north basin of a freshwater lake, Lake Biwa, Japan (35.13.2152 N, 135.59.7862 E). The collected sample was resuspended in Page’s amoeba saline, and large particles were removed by filtration. *A. castellanii* (Douglas) Page, strain Neff (ATCC 30010) was cultured in peptone-yeast extract-glucose (PYG) medium and seeded in a 96-well plate. The filtrated sample was inoculated into the amoeba culture with PYG medium supplemented with an antibiotic mix. A well that showed a cytopathic effect was collected and purified by the end-point dilution method. The presence of viruses was confirmed by negative staining as described previously (Fig. 1A) (Hikida et al. 2023).

### Genomic DNA extraction

The purified viruses were cultured in a 10-cm cell culture dish with the amoeba culture, and the supernatant was collected. The cultured viruses were collected by centrifugation at 9,000 rpm for 1 hour at 4°C (Sorvall ST8FR, Thermo Fisher Scientific) and resuspended in Tris-EDTA buffer (pH 8.0). The collected viruses were incubated overnight with 2 mg/mL of proteinase K and 1% sodium dodecyl sulfate. DNA was extracted by treatment with phenol twice and chloroform twice, followed by ethanol precipitation.

### Genome sequencing

Genomic DNA concentration was measured using a Qubit 4 fluorometer using Qubit dsDNA BR Assay Kits (Invitrogen). A sequencing library for long-read sequencing was prepared from 504 ng of the genomic DNA using the Ligation Sequencing Kit (SQK-LSK109, Oxford Nanopore Technologies), following the manufacturer’s protocol. The library was sequenced using an R9.4.1 Flongle flow cell (FLO-FLG001, Oxford Nanopore Technologies) on a MinION Mk1C with MinKNOW v22.05.08 software (Oxford Nanopore Technologies). A total of 291.2 Mb of data was obtained. The genomic DNA was also sequenced using the Illumina NovaSeq 6000 system in 150-bp paired-end mode. A total of 1 Gb of data was obtained. Quality control of the genomic DNA, library preparation, and sequencing were performed by Rhelixa, Inc. (Japan).

### Genome assembly

The raw long reads were assembled into contigs using miniasm v0.3 and minipolish v0.1.3 with default parameters (Li 2016). Source organisms for the assembled contigs were predicted by BLASTN implemented in BLAST+ v2.14.0 (Camacho et al. 2009) against the National Center for Biotechnology Information (NCBI) Nucleotide collection (nt/nr) database. Contigs likely derived from host organisms were removed. Based on long-read mapping to the assembled contigs using Minimap2 v2.22, the first assembly included an approximately 8-kbp region at the 5’ end of the putative genomic contig that showed higher coverage than the other region. In addition, no long reads connected this 8-kbp region to the putative viral genomic region. A BLASTN search revealed that the 5′ region likely originated from bacteriophage contamination. Therefore, we removed the raw long reads mapped to this phage-like region using Samtools v1.14 (Li et al. 2009). Reads mapped to the putative host contigs were also removed at this step. The remaining long reads were used for assembly as described above. Two contigs were constructed: one contig with 1,969,476 bp in length was typical for pandoravirus genomes, and the other 6,833 bp contigs were likely to be a part of the host mitochondrial genome based on a BLASTX search. Thus, we considered the 1,969,476-bp contig as the genomic sequence of the isolated virus. The Illumina short reads were mapped to this genomic contig by BWA v0.7.17 (Li 2013) and converted to a BAM format using Samtools, from which the sequence was polished with short reads using Pilon v1.23 (Walker et al. 2014).

The possibility of artifacts in the assembly process was examined by mapping the long reads used to assemble PanV-jap and PanV-biw to the PanV-biw genome. The long reads used for the PanV-jap assembly were retrieved from the DDBJ Sequence Read Archive. Mapping was performed using Minimap2, and the resulting SAM files were converted and sorted by Samtools. The mapping results were visualized by custom Python scripts.

### Genome annotation

Coding sequences in the genomic sequence were predicted by Prodigal v2.6.3 (Hyatt et al. 2010) with default parameters. The predicted amino acid sequences were further annotated by top hits from a BLASTP search against the NCBI non-redundant protein sequence (nr) database at the E-value threshold of 10^−4^ (Table. S1), followed by manual curation. tRNA was annotated using tRNAscan-SE v2.0.12 (Chan and Lowe 2019).

### Comparative genomics

The genomic sequences of the isolated pandoraviruses were retrieved from the NCBI Virus database (Table S2). ANI was calculated using fastANI v1.33 (Jain et al. 2018). Genomic regions similar to the newly isolated pandoravirus were identified by BLASTN search. Calculation of the GC ratio of each genome and visualization of the genome-wide comparison (e.g., dot plots) were performed using custom Python scripts.

### Phylogenetic analysis

Marker genes used for phylogenetic analysis of nucleocytoviruses were searched against coding sequences of the examined pandoraviruses using those encoded in PanV-sal as queries (Philippe et al. 2013). Because some genes were split into fragments, probably due to the insertion of introns or merely sequencing error, we selected RNA polymerase subunit B and AA18 helicase for inferring species trees. The protein sequences of these genes were aligned using MAFFT v7.520 (Katoh and Standley 2013) with the “auto” option. Phylogenetic trees were reconstructed using IQ-TREE 2 v 2.2.2.6 (Minh et al. 2020) with 1,000 replicates of the ultrafast bootstrap and Simodaira-Hasegawa approximate likelihood ratio test. Substitution models were determined using ModelFinder (Kalyaanamoorthy et al. 2017). The reconstructed trees were visualized using the iTOL web server (Letunic and Bork 2024).

Orthologous groups were identified from the pandoravirus genomes listed in Table S2 using Orthofinder v2.5.5 (Emms and Kelly 2019) with default parameters. In some pandoravirus genomic sequences, the protein-coding sequences are not available. For these genomes, we predicted protein-coding genes de novo using Prodigal with default parameters. Protein sequences belonging to an OG of morn-repeat domain proteins were aligned, and a phylogenetic tree was built and visualized, as described above.

## Data access

The short- and long-read sequencing data were deposited in DDBJ with the accession numbers DRR641321 and DRR641322, respectively. The genomic sequence was deposited in DDBJ with the accession number LC870878.

## Acknowledgments

Field sampling was supported by the Center for Ecological Research, Kyoto University, a Joint Usage/Research Center. The electron microscopy study was supported by the Division of Electron Microscopic Study, Center for Anatomical Studies, Graduate School of Medicine, Kyoto University. Computation time was provided by the SuperComputer System, Institute for Chemical Research, Kyoto University. This study was supported by the Japan Society for the Promotion of Science KAKENHI grant numbers 22K15175, 21J00174 to HH, and 22H00384 to HO.

## Competing interest statement

The authors declare no competing interests.

## Notes

### Competing Interest Statement

The authors have declared no competing interest.

