## Supplementary material for "Horizontal transfer of a 180-kbp genomic fraction among the largest viral genomes": Table S2, Fig. S1, Fig. S2, Fig. S3, Fig. S4, Fig. S5

### Horizontal transfer of a large genomic fraction between giant pandoraviruses.

**Table S1 (Separate Excel file)**
**Table S2**
**Figure S1–S5**

‡Present address: Department of Virology I, National Institute of Infectious Diseases, Japan Institute for Health Security, 1-23-1, Toyama, Shinjuku, Tokyo 162-8640, Japan; Research Center for Biosafety, Laboratory Animal and Pathogen Bank, National Institute of Infectious Diseases, Japan Institute for Health Security, 1-23-1, Toyama, Shinjuku, Tokyo 162-8640, Japan

**Table S2. List of viral genomes used in this study**

| Virus name* | Accesssion | Abbreviation | Clade |
|---|---|---|---|
| Pandoravirus neocaledonia | MG011690.1 | PanV_neo | Clade B |
| Pandoravirus aubagnensis | MZ420563.1 | PanV_aub | Clade B |
| Pandoravirus japonicus | LC625835.1** | PanV_jap | Clade A-I |
| Pandoravirus pampulha strain Biwa | LC870878 | PanV_biw | Clade A-I |
| Pandoravirus lena* | OQ411594.1 | PanV_len | Clade A-II |
| Pandoravirus lena* | OQ411595.1 | PanV_len | Clade A-II |
| Pandoravirus lena* | OQ411596.1 | PanV_len | Clade A-II |
| Pandoravirus lena* | OQ411597.1 | PanV_len | Clade A-II |
| Pandoravirus lena* | OQ411598.1 | PanV_len | Clade A-II |
| Pandoravirus lena* | OQ411599.1 | PanV_len | Clade A-II |
| Pandoravirus pampulha strain 8.5 | LT972219.1 | PanV_pam | Clade A-I |
| Pandoravirus salinus | KC977571.1 | PanV_sal | Clade A-II |
| Pandoravirus mammoth* | OQ411600.1 | PanV_mam | Clade B |
| Pandoravirus mammoth* | OQ411601.1 | PanV_mam | Clade B |
| Pandoravirus dulcis | KC977570.1 | PanV_dul | Clade A-I |
| Pandoravirus talik | OQ413801.1 | PanV_tal | Clade A-II |
| Pandoravirus massiliensis isolate BZ81 c | MZ384240.1 | PanV_mas | Clade B |
| Pandoravirus celtis | MK174290.1 | PanV_cel | Clade A-II |
| Pandoravirus kuranda isolate Kuranda | ON887157.1 | PanV_kur | Clade B |
| Pandoravirus inopinatum isolate KlaHel | KP136319.1 | PanV_ino | Clade A-II |
| Pandoravirus belohorizontensis | MZ420562.1 | PanV_bel | Clade A-I |
| Pandoravirus quercus | MG011689.1 | PanV_que | Clade A-II |
| Pandoravirus macleodensis | MG011691.1 | PanV_mac | Clade B |

*Contigs are registered separately.

**re-asemmbled in this study

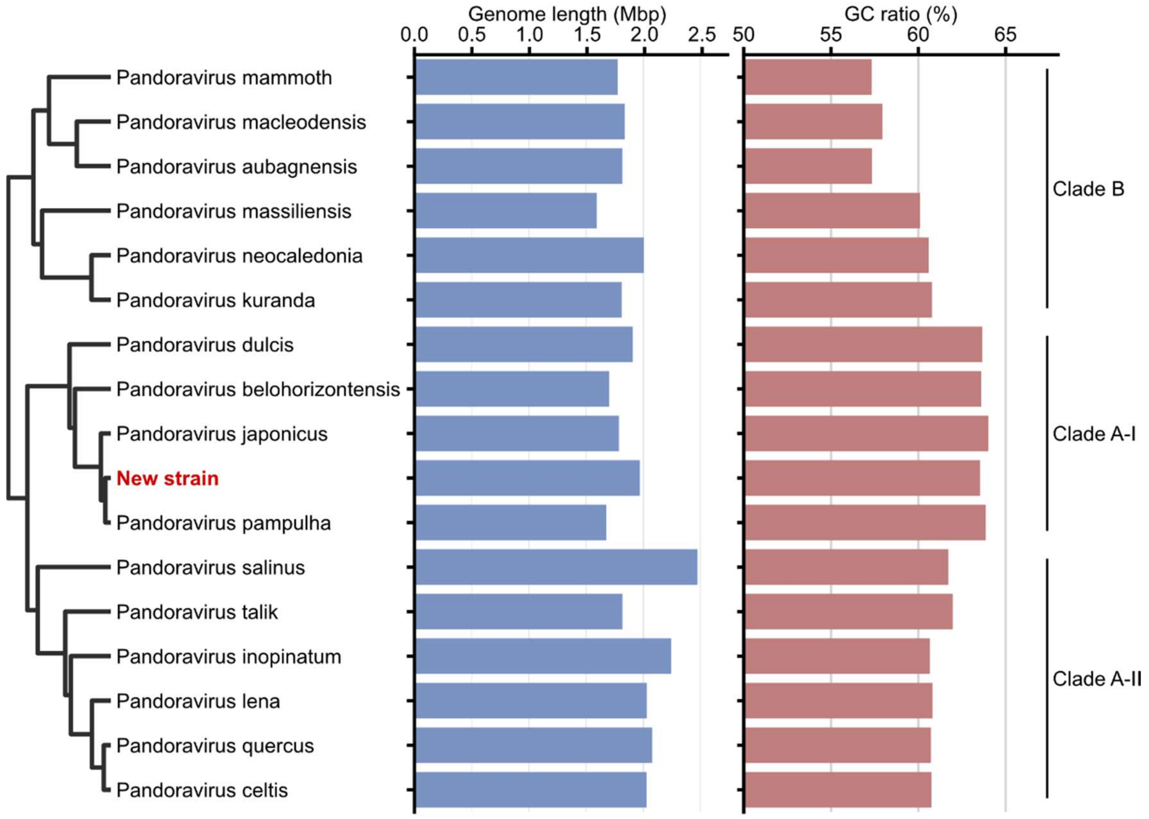


**Figure S1. Genomic properties of pandoraviruses.**

Length and GC ratio of pandoravirus genomes. The left dendrogram represents clustering based on the average nucleotide identity shown in Figure 1C.

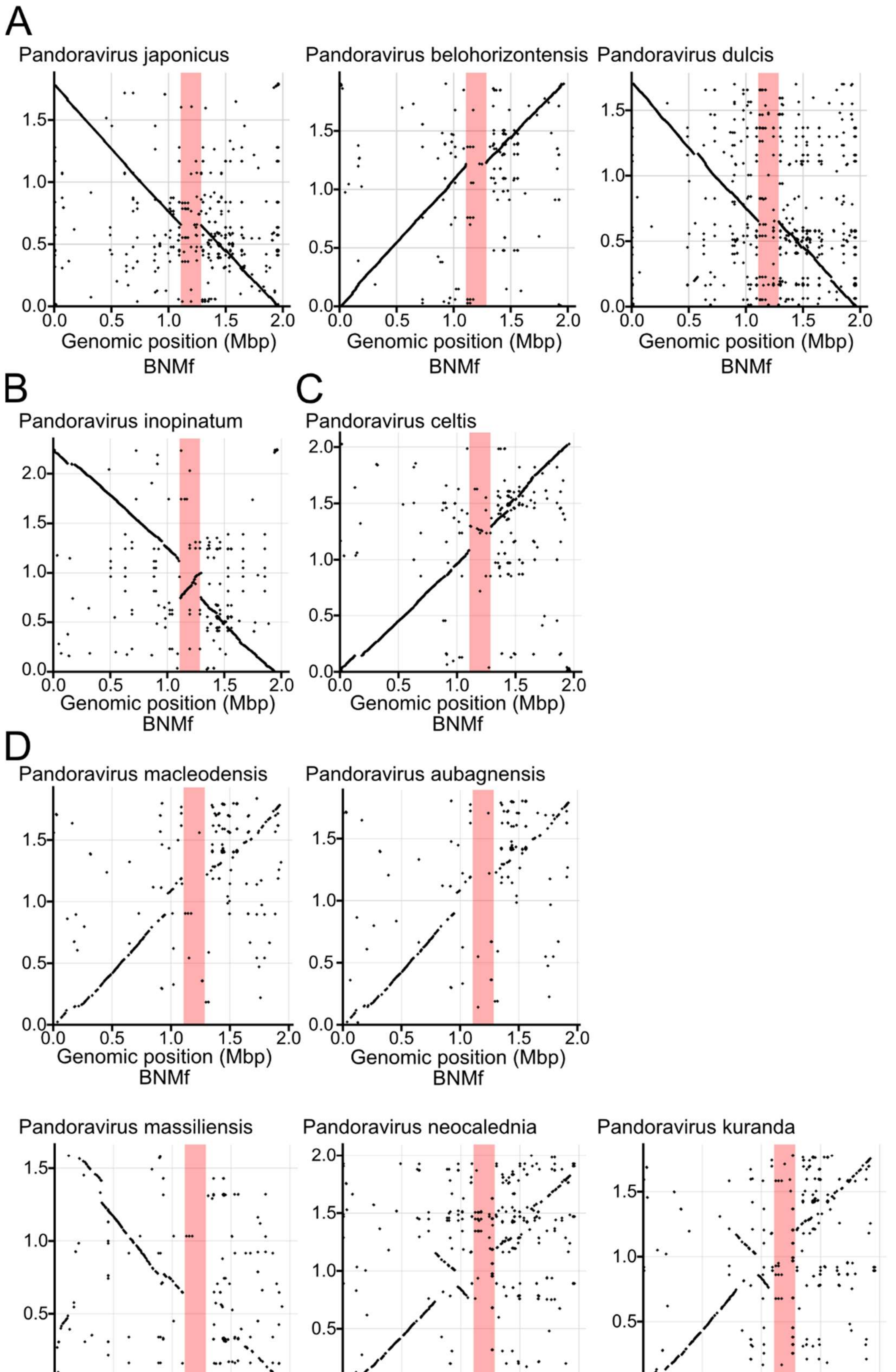
A
Pandoravirus japonicus
Pandoravirus belohorizontensis
Pandoravirus dulcis
B
Pandoravirus inopinatum
C
Pandoravirus celtis
D
Pandoravirus macleodensis
Pandoravirus aubagnensis
Pandoravirus massiliensis
Pandoravirus neocalednia
Pandoravirus kuranda
Genomic position (Mbp)
BNMf

**Figure S2. Genomic comparison between PanV-biw and the other pandoraviruses.**

Genomic comparison between PanV-biw and (A) pandoraviruses belonging to Clade A-I, (B, C) Clade A-II, and (D) Clade B. The X- and Y-axes represent the genomic position in PanV-biw, and other viruses, respectively. The viruses are designated at the top left. Red boxes indicate the 180-kbp region missing in other Clade A-I viruses.

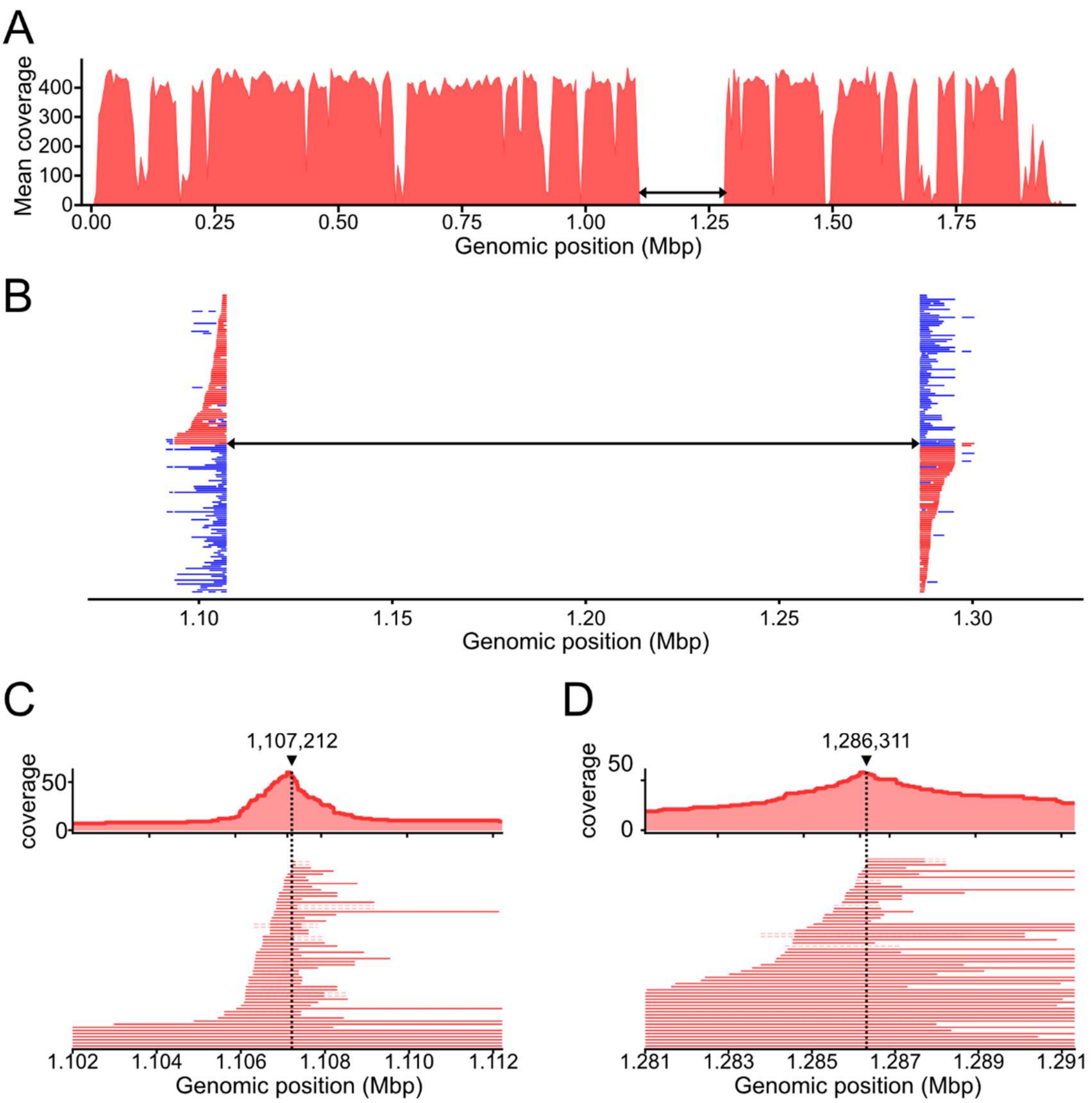


**Figure S3. Read-mapping results for PanV-jap and PanV-biw.**

(A) Raw long reads from PanV-jap were mapped to the PanV-biw genome. (B) Enlarged image around the 180-kbp region in (A). Each row represents a long read that bridged the 5′ and 3′ flanking regions of the 180-kbp region. The primary and secondary alignments of each read are shown in red and blue, respectively. (A, B) The black arrow indicates the 180-kbp region missing in PanV-jap. (C, D) Raw long reads of PanV-biw were mapped to the PanV-biw genome. Only reads bridging the 180-kbp region and its flanking region were shown. (C) 5′ and (D) 3′ boundary between the 180-kbp and flanking regions.

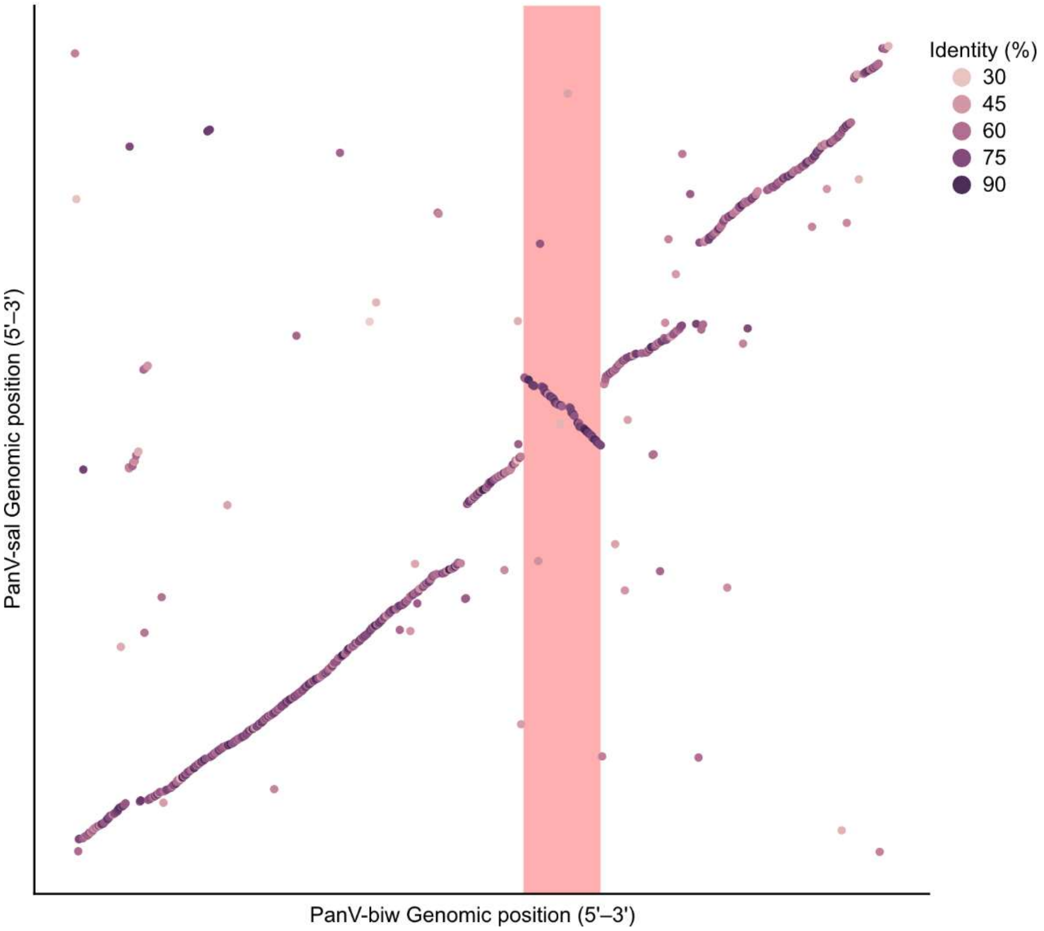


**Figure S4. Consistent synteny between PanV-biw and PanV-sal.**

Protein-level synteny between PanV-biw and PanV-sal was visualized from the data used in Figure 3A. The X- and Y-axes indicate the genomic positions of PanV-biw and PanV-sal, respectively. The colors indicate the identity of the proteins. Red boxes indicate the 180-kbp region in PanV-biw.

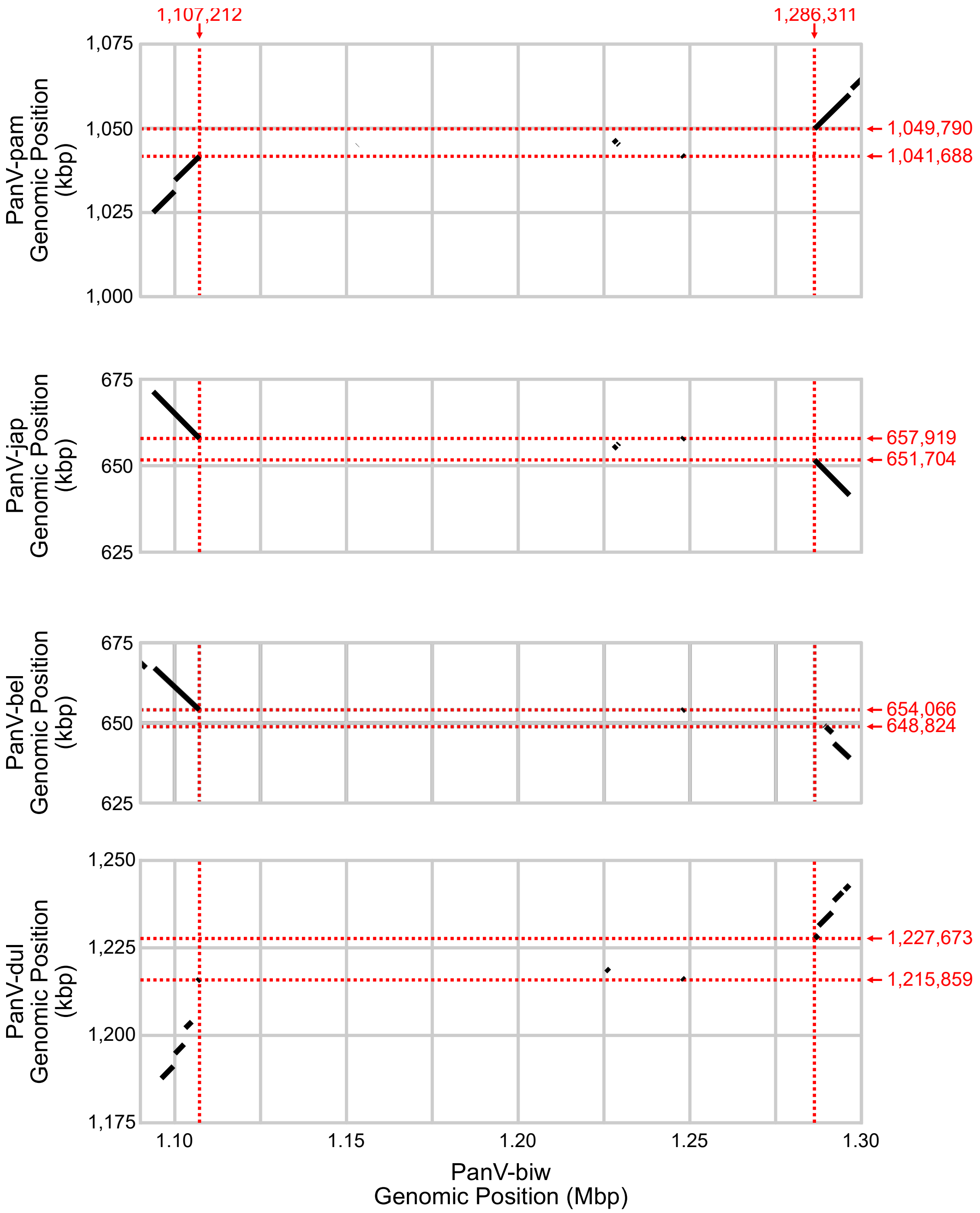


**Figure S5. Enlarged dot plots around the regions corresponding to the 180-kbp region in PanV-biw**

Genomic comparison between PanV-biw and pandoraviruses belonging to Clade A-I. Flanking regions of the 180-kbp region were enlarged. X- and Y-axes represent the genomic position in PanV-biw and Clade A-I pandoraviruses, respectively. The viruses are designated on the left. The vertical and horizontal red dotted lines indicate the manually determined boundaries of the 180-kbp region and the corresponding regions of each Clade A-I pandoraviruses.
